# Past, present, and future spatial distributions of deep-sea coral and sponge microbiomes revealed by predictive models

**DOI:** 10.1101/2024.03.04.583289

**Authors:** Kathrin Busch, Francisco Javier Murillo, Camille Lirette, Zeliang Wang, Ellen Kenchington

**Author notes:** **Competing Interests statement** The authors declare no competing interests.

## Abstract

Knowledge of the spatial distribution patterns of biodiversity in the ocean is key to evaluate and ensure ocean integrity and resilience. Especially for the deep ocean, where *in situ* monitoring requires sophisticated instruments and considerable financial investments, modelling approaches are crucial to move from scattered data points to predictive continuous maps. Those modelling approaches are commonly run on the macrobial level, but spatio-temporal predictions of host-associated microbiomes are not being targeted. This is especially problematic as previous research has highlighted that host-associated microbes (microbiomes) may display distribution patterns that are not perfectly correlated with host animal biogeographies, but also with other factors such as prevailing environmental conditions. We here establish a new simulation approach and present predicted spatio-temporal distribution patterns of deep-sea sponge and coral microbiomes, making use of a combination of environmental data, host data and microbiome data to advance our understanding of deep-sea microbiomes. This approach allows predictions of microbiome spatio-temporal distribution patterns on scales that are currently not covered by classical sampling approaches at sea. This includes both predictions in space within regional oceanic provinces off eastern North America, and also in time, with predictions into the past and future, covering a time span of 214 years. In summary, our presented predictions allow (i) identification of microbial biodiversity hotspots in the past, present, and future, (ii) evaluation of microbial-macrobial connections at case-study sites through trait-based predictions, and (iii) identification of shifts in microbial community composition (key taxa) across environmental gradients and shifting environmental conditions.

## Introduction

Assessments and predictions of spatial distribution patterns have been conducted extensively for several deep-sea sponge (^1;2^) and deep-sea coral species (^3;4;5;6;7;8;9^). Although microbial (*16S* amplicon) data exist and the main driving factors have been identified for both deep-sea sponge (^10^;^11^) and coral (^12–14^) microbiomes, those existing datasets represent only snapshots in space and time, and microbial information have not been incorporated into host distribution models. We consider the generation of continuous tempo-spatial distribution patterns as a next major research direction to answer research questions such as: “Where may biodiversity hotspots of host-associated microbiomes be found in the contemporary and future ocean?”, and subsequently, “How do predicted shifts in the future align with predicted patterns of the past?”. Additional questions are: “How do host-associated microbial communities shift across environmental gradients and how do abundances of key microbial taxa vary spatially?”, and lastly “How do patterns of host-associated microbial abundances relate to spatial niche separations and distribution patterns of the larger reef community?”. Our study aim for this study was to answer those questions for deep water sponge and coral-associated microbiomes, as both, sponges as well as corals represent key animal species of deep ocean benthic communities.

## Results & Discussion

Using five key species of North American east coast waters, *Weberella bursa, Stryphnus fortis, Lophelia pertusa* [*Desmophyllum pertusum*], *Desmophyllum dianthus*, and *Vazella pourtalesii*, coral- and sponge-associated microbial biodiversity hotspots were revealed in the eastern Canadian Arctic, the Flemish Cap, the Scotian Shelf, and in US waters from the border with Canada along the Eastern Seaboard to Florida (**Figure 1A**). The underlying newly established simulation approach was to model the host distributions first (**Supplementary Figure 1, Supplementary Table 1**), using *in situ* measurements and nine explanatory environmental parameters, and to then overlay the measured host species-specific microbiome data. A similar approach was used to make predictions under past and future environmental conditions. Strongest shifts under future projections (mainly gains in cumulative microbial richness) were predicted to occur in the Gulf of Maine and the Laurentian Channel, while losses in cumulative microbial richness in coral and sponges were predicted to occur in the Canadian Arctic. Overall, the fraction of grid cells with observed losses in coral and sponge-associated cumulative microbial richness was predicted to increase in the future in comparison to the past (**Figure 1B**). This highlights that host-associated microbial landscapes are highly dynamic in time, and that spatial areas and ecosystems have faced continuous invasions and introductions of new metaorganisms (host + microbiome) over the 214 year time frame analysed.

**Figure 1.**
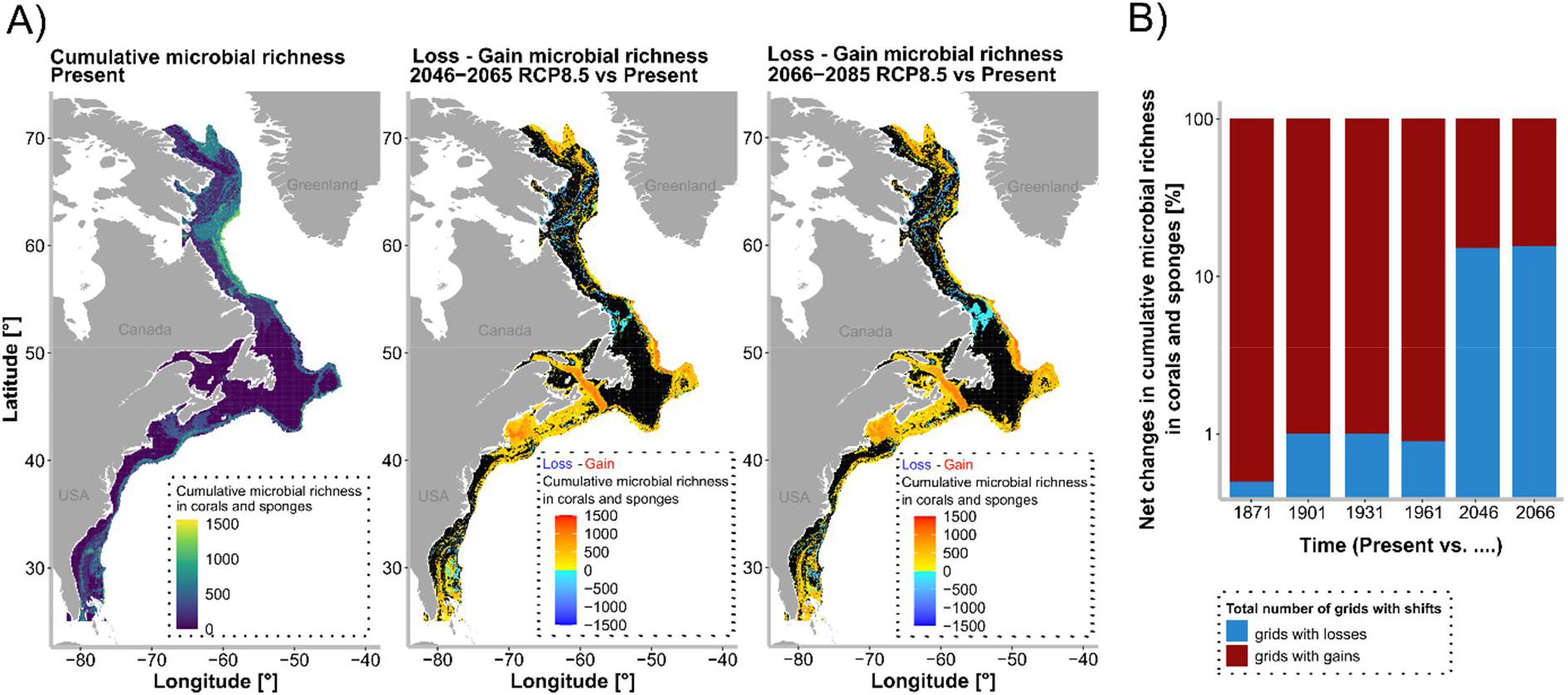
Microbial biodiversity hotspots in the past, present, and future. **A)** Predictions of cumulative microbial richness in deep-sea corals and sponges for the present, as well as predicted gains and losses in cumulative microbial richness for the two future timeframes. **B)** Net changes in cumulative microbial richness in corals and sponges over time, given as percentages of total shifted area.

Due to the observed habitat shifts over time for sponges and corals, we aimed to dig deeper into modelling spatial distribution patterns of individual microbial taxa in relationship to gradients and shifts in environmental parameters. The *Vazella pourtalesii* microbiome was chosen as a study system for this approach, and we focussed only on “region B” (which includes the shelf off Nova Scotia; part of the Gulf of Maine, and areas off Newfoundland; for more details see **Methods** and **Supplementary Material**) for this analysis because shifts in cumulative microbial richness were predicted to be largest in this region for the two future timeframes (**Supplementary Figure 2**). Two key microbial taxa (ASVs) with a high relative abundance and prevalence, as well as a strong temperature-relationship were picked (for more details on key microbial taxa see **Supplementary Text 1**, and **Methods**). In order to detect shifts of those taxa across a very fine/continuous temperature gradient (which can also guide the design of future lab experiments), microbial key community compositions were predicted across a fine-scale temperature gradient based on a HMSC-modelling approach. We identified different optimal temperatures for the two key microbial taxa at 7.5 °C for ASV1 and 7.7 °C for ASV2 (for a temperature range between 6.2-8.2°C), and detected slight differences in their response to changes in temperature (**Figure 2A**). We focused on temperature as it has repeatedly been identified as a major driving factor of sponge microbial community composition (e.g. ^10,15^). Predicted spatial distribution maps (**Figure 2B**) were then generated for the two temperature-related key microbial taxa, providing for the first time a continuous picture of how relative abundances of *Vazella pourtalesii*-associated microbial taxa may vary geographically. But how do changes in key microbial taxa translate into overall microbial community composition? In order to evaluate how shifts in relative abundances of the key temperature-driven ASVs translate into shifts in overall microbial community composition, we constructed co-occurrence networks. Those co-occurrence networks suggest that changes in abiotic conditions (temperature) may not only impact single key microbial taxa, but may trigger cascades through a modular network of microbial taxa (**Supplementary Text 2, Supplementary Figure 3**) .

**Figure 2.**
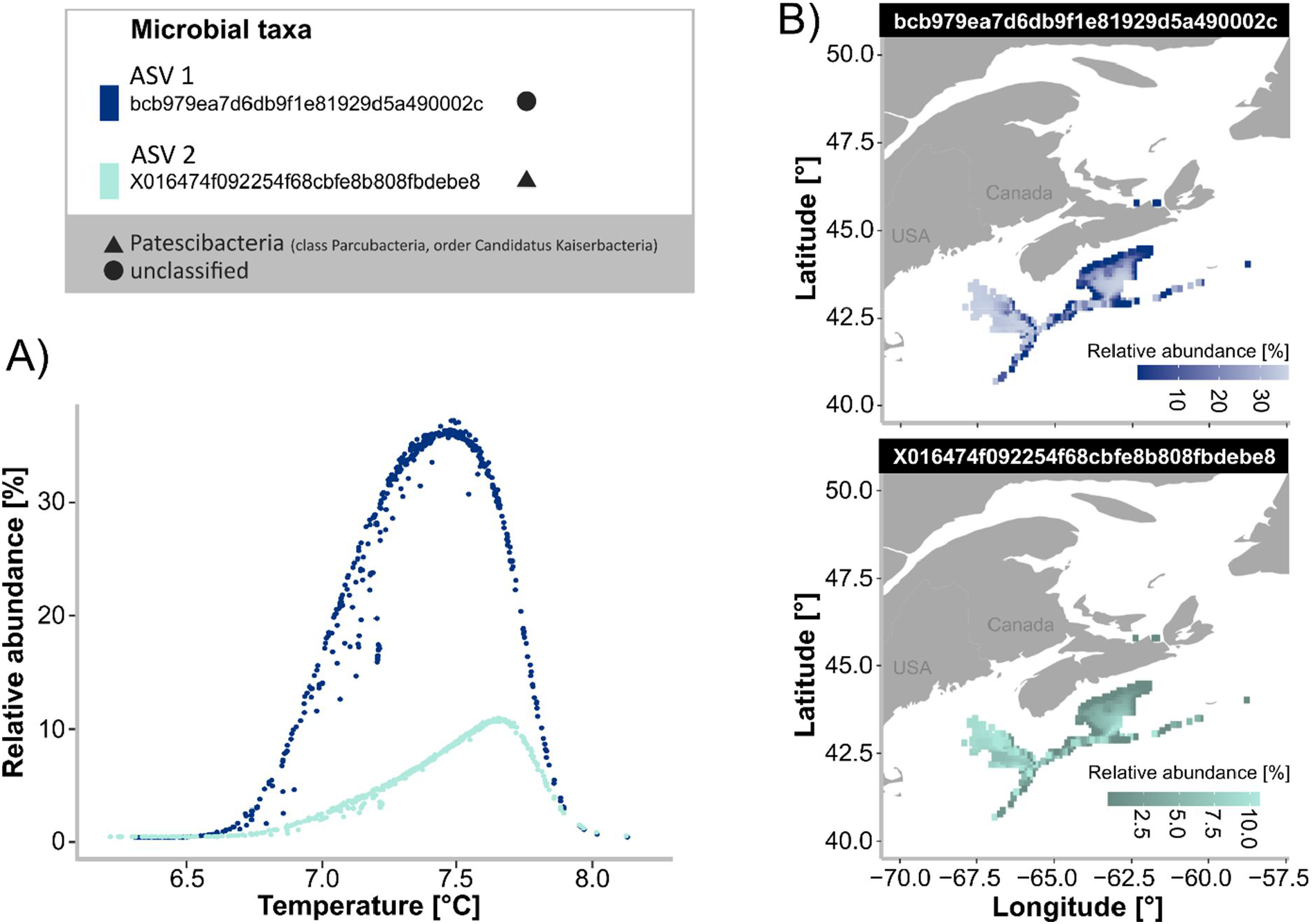
Relative abundances of key microbial taxa across shifting environmental conditions and environmental gradients. Two temperature-related key ASVs are shown by different colors, their taxonomic identity is indicated by a triangle (Patescibacteria (class Parcubacteria, order Candidatus Kaiserbacteria)), and circle (unclassified). **A)** Predicted relative abundances of key ASVs with good model fit in the glass sponge *Vazella pourtalesii* over a fine-scale temperature gradient. **B)** Predicted distribution maps of relative abundances of key ASVs with good model fit in the glass sponge *Vazella pourtalesii*.

In summary, this study identified microbial biodiversity hotspots in the past, present, and future along the North American east coast, pinpointed temperature-related shifts in microbial community composition, and derived individual geographic maps showing relative abundances of key microbes across an environmental gradient. In addition we also tested the usability of trait-based modelling approaches to improve our holistic understanding of (deep-sea) reef ecosystems. The underlying idea here was to assess potential niche separations of sponge holobionts based on their microbial abundance status. We found a spatial niche separation between HMA (high microbial abundance) and LMA (low microbial abundance) sponges and also different co-occurring sessile filter feeding invertebrates with each of the two sponge types (so presumably different associated food webs). We thus believe that this trait-based approach is helpful to assess biodiversity-ecosystem function relationships and microbial-macrobial interactions, and we provide more information and results in the **Supplementary Material** (**Supplementary Text 3, Supplementary Figure 4**). Taken together, all our results suggest that, despite a cumulative uncertainty (added up for each modelled level of organisation: the environmental predictors, the host distributions, the microbiome predictions), spatio-temporal predictions of host-associated microbial communities can reveal interesting ecological patterns, help to fill data gaps in undersampled habitats, and may guide future experiments.

## Acknowledgements

This work was supported by a fellowship of the German Academic Exchange Service (DAAD) to KB (91855556). Further funding was through Fisheries and Oceans Canada, Maritimes Region project “Mapping Biodiversity and Ecosystem Services of Benthic Communities on the Scotian Shelf”, Marine Conservation Targets Project ID: 395, to EK and FJM. We appreciate a helpful discussion with Christina Kellogg about deep-sea coral microbiomes.

## Data availability statement

The data and data links of this paper were deposited in the Mendeley database - reserved doi:10.17632/fx3vd2tgcf.1.

## Methods

The applied workflow is described in the following. The individual analysis steps, were executed in R (version 4.1.3; ^1^), ArcGIS Pro (version 2.8.8; ^2^), and Python (version 3.7.3; ^3^).

### Spatial extent and preparation of environmental data layers for species distribution modeling

The spatial extent of this study covered an area off the North American east coast in the northwest Atlantic, spanning a large latitudinal extent from ∼25°N (Nassau, Bahamas) to ∼71°N (northern part of Baffin Island, Canada), (**Supplementary Figure 5B**). In order to avoid inclusion of land points, a 5 km land buffer was applied along the coastline, and all environmental grid cells falling within this boundary were removed. The area extended seaward from the buffered coastline to the 2000 m bathymetric contour; any areas deeper than 2000 m were excluded from the study area. Raster cell size was 0.088°, using the WGS 1984 datum. With the ‘Calculate Geometry’ tool in ArcGIS Pro the geodesic area of each grid cell was calculated, showing that grid cells at the North extended to approximately 31km^2^, grid cells at the south extended to around 87km^2^, and grid cells at the middle coordinate of the whole study area (∼48 °N) were around 64 km^2^. The study region was broken up into three different regions (A, B, C, **Supplementary Figure 5B**) in order to perform analysis on a sub-regional level and to allow comparability to previous studies (e.g. ^4^). Region A, the northernmost subregion, spanned a geographic range from 52.1° N to 71.5°N; while region C, the southernmost subregion, spanned a geographic range from 25.0° to 40.0°. Region B was the subregion in between region A and region B.

Time-wise this study spans 214 years, from 1871 until 2085. Two static (stable in time) environmental parameters were used throughout the complete time span. These were “bottom depth” (derived from the General Bathymetric Chart of the Oceans (version 2019; ^5^), and “slope”. Slope, in degrees, was derived from the bathymetry raster using the Slope Tool in ArcGIS Pro’s Spatial Analyst toolbox, after projecting the bathymetry layer to Albers projection for this analysis (and then back again to WGS 84). Bilinear interpolation was used to resample both depth and slope to match the resolution of the dynamic climatic variables (0.088°).

Dynamic (varying in time) environmental layers for our study area were derived from two sources: the Bedford Institute of Oceanography North Atlantic Model (BNAM; ^6^), and the Simple Ocean Data Assimilation reanalysis of ocean climate variability (SODA; ^7^). The complete study period was split into 7 time frames covering the years 1871-1900, 1901-1930, 1931-1960, 1961-1989, 1990-2015, 2046-2065, and 2066-2085 (**Supplementary Figure 6**). The two future time frames were simulated under IPCC^8^ worst-case CO_2_ emission scenarios (RCP8.5; ^9^). As future time frames of BNAM were provided in delta fields, absolute values of each variable were calculated for inputting them into the species distribution models. All data derived from BNAM were calculated as averages across the complete time period of each multi-annual timeframe. Dynamic environmental layers for the present and future were mean bottom temperature, mean bottom salinity, mean bottom current velocity, mean bottom stress, mean maximum mixed layer depth, mean surface temperature, and mean surface salinity (**Supplementary Figure 7**). Mean bottom current velocity (U_b_) was calculated from eastward seawater velocity (U) and northward seawater velocity (V), with the following formula: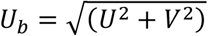. BNAM (and BNAM-derived) point data were used to create continuous raster surfaces in ArcGIS Pro (Point to Raster tool). All layers were displayed using WGS 84 geographic coordinates and the final raster resolution was 0.088°.

For the past, data were derived from the Carton-Giese SODA 2.2.4, 1871-2008 Assimilation Run ^10^, in a monthly-averaged form. Overall averages across each of the roughly 30-years-timeframes were calculated. Bottom layers were calculated for mean temperature, mean salinity, mean u, and mean v with the help of the GEBCO bathymetry layer of the study area. In particular, the respective value was chosen from the SODA depth layer that was closest to the bottom depth at each grid coordinate. Mean bottom current velocity was calculated from mean U and mean V using the same formula as stated above for the BNAM data. Furthermore, using the mean bottom layer *U*_*b*_, the strength of the bottom stress (*τ*_*b*_) was calculated as *τ*_*b*_ = 3.5 × 10^-3^ × *ρ* × *U*_*b*_^2^ for BNAM, and as 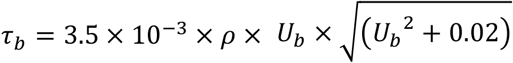 for SODA; where, p is the density of seawater [kg m^-3^]. The density variable needed for the previous calculation was calculated with the help of the *gsw*-package in R as follows: In the first step pressure was calculated with the help of the *gsw_p_from_z* function, using depth and latitude as input variables. In the second step, absolute salinity (SA; g kg^-1^) was calculated from mean bottom salinity [psu] and pressure [dbar] (using the function *gsw_SA_from_SP*). In the third step conservative temperature (CT) was calculated from mean bottom temperature [°C], absolute salinity [g kg^-1^], and pressure [dbar] (using the function *gsw_CT_from_t*). In the final step, density was calculated from absolute salinity [g kg^-1^], conservative temperature [°C], and pressure [dbar] (using the function *gsw_rho*). Maximum mixed layer depths was calculated for each grid coordinate using the *castr*-package in R (in particular the *mld*-function), after extracting vertical depth-profiles of temperature and salinity over the complete water column at each grid coordinate, and calculating vertical density profiles following the density-calculation procedure described above. The 5 m-depth layer of SODA was taken to represent surface temperature and mean surface salinity. For all SODA layers, point data was interpolated using ordinary kriging in ArcGIS Pro, in order to match the spatial resolution (0.088°) of the BNAM layers. As the SODA data did not cover parts of the Hudson Strait/Ungava Bay, NA values were added to this area covered by BNAM in order to have the same spatial extent for both datasets. An overview containing all used environmental data layers was archived *doi:10*.*17632/fx3vd2tgcf*.*1 (“Data2”)*.

### Coral and sponge occurrence data

The aim of this study was to cover key coral and sponge species of the study area, which occur in high abundances, and for which information on the associated microbiome was available: *Weberella bursa, Stryphnus fortis, Lophelia pertusa, Desmophyllum dianthus*, and *Vazella pourtalesii* were found to match those criteria. A total of 11,931 raw coral and sponge occurrences were compiled for the study area. Occurrence records of *Weberella bursa, Stryphnus fortis, Lophelia pertusa, Desmophyllum dianthus*, and *Vazella pourtalesii* were drawn from multiple published sources (^4;11;12;13/14;15;16;17;18^), as well as from previously unpublished data. For a more detailed overview of the individual data records used in this study, see *doi:10*.*17632/fx3vd2tgcf*.*1 (“Data3”)*. Total raw occurrence records compiled from these sources, were as follows: 925 *Weberella bursa*-related individual records (110 presences and 815 absences), 694 *Stryphnus fortis*-related individual records (69 presences, 625 absences), 2,103 *Lophelia pertusa* presence records, 3,629 *Desmophyllum dianthus* presence records, and 4,580 *Vazella pourtalesii*-related records (164 presences, 4,416 absences). For each coral or sponge species, raw occurrence records were filtered prior to modelling, such that only one record per grid cell was retained, and coordinates of the grid cell’s centroid were assigned to the record. If presences and absences occurred in the same grid cell, presence was used. Mapping to grid cells resulted in 99 cells with *Weberella bursa* presences, 55 cells with *Stryphnus fortis* presences, 142 cells with *Lophelia pertusa* presences, 84 cells with *Desmophyllum dianthus* presences, and 130 cells with *Vazella pourtalesii* presences (see *doi:10*.*17632/fx3vd2tgcf*.*1 (“Data4”)*). Cells which had recorded absences were filled with zeros. In addition, pseudo-absences were added for each species to those cells which had recorded presence-absences for any of the other coral or sponge species, assuming that the respective species would have been recorded if present. Following this procedure, all five coral and sponge species had records (presence + absence) in 5477 grid cells of our study area (i.e. there were records in ∼15 % of all grid cells).

### Random Forest predictions of five coral and sponge species occurrence

In order to predict coral and sponge distribution patterns, Random Forest (RF) models were fitted using the *randomForest*-package in R ^19^ with default parameters and 500 trees. Model accuracy metrics and threshold probabilities were derived using 5-fold spatial block cross-validation. For each cross-validation run the threshold-independent area under the receiver operating characteristic curve (AUC) was calculated, and the mean and standard deviation were derived. The *optimal*.*thresholds*-function in the “PresenceAbsence”-R package ^20^ was used to calculate several common threshold values above which a given relative probability of occurrence is considered a presence. The threshold which maximizes the sum of sensitivity and specificity (MSS) was our threshold of choice. There, sensitivity and specificity represent the proportion of accurately predicted presences and absences, respectively. Using MSS, probabilities of occurrence outcomes from each cross-validation run were converted into predicted binary outcomes which were subsequently summarized into a 2x2 confusion matrix. In order to assess model performance along with AUC, the sensitivity, specificity, and the true skill statistic (TSS; ^21^) were derived. The *importance*-function in the R package “randomForest” was applied to evaluate the importance of the environmental predictor variables in the RF models using the mean decrease in Gini index. Functional response curves were produced to assess changes in the relative probability of occurrence across environmental variable gradients (after ^22^) (**Supplementary Figure 8**).

### Computation of cumulative microbial richness

In total 99 samples were processed to analyse microbiome diversity (*16S* amplicon data). These 99 samples consisted of 16 *Weberella bursa*, 19 *Stryphnus fortis*, 8 *Lophelia pertusa*, 7 *Desmophyllum dianthus*, and 32 *Vazella pourtalesii* individuals (**Supplementary Figure 5A**). The raw sequences were retrieved from the European Nucleotide Archive (ENA) database and originate from ^11;12;17^ (for details on accession numbers see *doi:10*.*17632/fx3vd2tgcf*.*1 (“Data5”)*). As the same region of the 16S gene was amplified for all samples (V3 region), all reads were processed together within the QIIME2 environment (version 2019.10; ^23^). Processing was done according to the bioinformatic pipeline published in ^11^. In the present study, primers and heterogeneity spaces were removed. Then additional 15nt were trimmed at the start of all reads, and the reads were truncated to a length of 225nt. Chimeric reads were removed from the dataset. Amplicon sequence variants (ASVs) were generated from forward reads with the DADA2 algorithm ^24^. The FastTree2 plugin was used to calculate phylogenetic trees based on resulting ASVs. Classification of representative ASVs was done with help of the Silva 132 99 % OTUs 16S database ^25^ with the help of a *16S* region-specific trained naïve Bayes taxonomic classifier. Afterwards, chloroplast, mitochondrial, eukaryotic, and unassigned reads were removed. A sampling depth of 6000 was applied, and alpha diversity indiced (Shannon indices) calculated for each of the 99 samples. Mean microbial richness estimates were calculated for all five coral and sponge species, and the seawater reference group. This revealed a mean number of observed ASVs of 161 ASVs in *Weberella bursa*, 480 ASVs in *Stryphnus fortis*, 149 ASVs in *Lophelia pertusa*, 428 ASVs in *Desmophyllum dianthus*, 336 ASVs in *Vazella pourtalesii* individuals, and 583 ASVs in seawater (see **Supplementary Figure 5C**). Cumulative microbial richness was calculated for each grid cell, by combining coral or sponge occurrence with microbial richness estimates. In particular, predictions of relative probability of animal occurrence were thresholded into a binary depiction of suitable vs unsuitable habitat using the individual MSS values used to threshold the confusion matrices of each animal species. Afterwards these binary predictions were overlaid for all five animal species, and for those animals with a presence in the respective grid cell, mean microbial richness estimates of the respective animal species were summed up (and called “cumulative microbial richness”).

In order to identify shifts in microbial richness over time (present vs. future), the relative probabilities of animal occurrence projected for the two future timeframes (2046-2065, and 2066-2085) were also thresholded and cumulative microbial richnesses were calculated using the same procedure as described above. Afterwards, cumulative microbial richnesses were compared for those three time frames (present and two future timeframes), but also for the four past timeframes. The “Calculate Geometry” tool in ArcGIS Pro was used to calculate the geodesic area of each grid cell [km^2^]. From there summed total areas [km^2^] were calculated of all grid cells per species for each year (**Supplementary Table 2**). Areas that experienced a gain, loss, or no change in cumulative microbial richness from present-day conditions were evaluated. In order to get a feeling for the magnitude of shifts across the complete study area, the overall area of shifts (including both, areas of gain and areas of loss) was calculated. For this, grid cells showing a shift over time were summed and thereof the percentages of gains and losses were calculated, respectively (**Supplementary Figure 2**).

### Hierarchical Modelling of Species Communities (HMSC) to predict abundance shifts of key microbes across environmental gradients

In order to predict abundance shifts of microbes along changing environmental conditions, we focused on the microbiome of the glass sponge *Vazella pourtalesii*, as microbial sample numbers were largest for this species (n=32). Based on the overall microbial community compositions, that were derived from 16S amplicon sequencing analyses (see above), “key microbial taxa” were calculated. In order to define key microbial taxa of the *Vazella pourtalesii* microbiome, the total microbial community was divided as follows: (i) “core ASVs” were defined as those ASVs occurring in all animals, (ii) “variable community ASVs above 10” are those ASVs occurring in ten or more animals, (iii) “variable community ASVs below 10” are those ASVs occurring in between two to nine animals, (iv) “individual ASVs” are those ASVs occurring in one animal only. In order to identify key microbial taxa, we consider ASVs that occur in ten or more *Vazella pourtalesii* individuals only, and which have a mean relative abundance of more than 1 %. Following this procedure, 15 key ASVs were identified, which cover 67 % of the total microbial community of *Vazella pourtalesii* in terms of relative abundance. After identification of key microbial taxa, our data was brought into the format needed to run the HMSC model: Relative abundances of the 15 key ASVs were standardized to 100 %. The bottom temperature raster and the predicted *Vazella pourtalesii* occurrence raster were merged into a new raster, containing coordinates and temperature values at locations of predicted *Vazella pourtalesii* occurrences only (i.e. presence probabilities above the respective MSS threshold identified for *Vazella pourtalesii*). The study area for running HMSC models was subsetted to study region B only, as this was the area with highest predicted area shifts in cumulative microbial richness (**Supplementary Figure 2**). In region B, *Vazella pourtalesii* is predicted to occur in a temperature range between 4.6°C and 9.2°C. Coordinates of *Vazella pourtalesii* individuals sampled *in situ*, for molecular work, were matched with the resulting temperature grid by finding the closest points of sampling coordinates in the temperature grid coordinates. Microbial communities of *Vazella pourtalesii* falling into the same grid cell were averaged. After doing so, four grid cells (locations) were used as input for the HMSC model. In order to enlarge the number of locations in the model, we extracted additional locations (coordinates) with the same temperature as the in situ measurements (precision of two digits °C) from the subsetted temperature grid. Our applied rational was that we expect a very similar mean microbial community composition in the same sponge species under the same temperature conditions. We thus calculated average relative microbial abundances of key ASVs per temperature category (i.e. the individual temperature values at all four sampling locations with a precision of two digits °C), and transformed them into “fake” integers for the HMSC modelling. We then appended the data of the 18 additional locations, identified as locations with similar temperatures to the *in situ* sampling locations (see description above), using microbiome averages of each respective temperature category as additional data values. This procedure lead to 22 locations in total, that were used in the HMSC model as the spatial training matrix of microbiome data. We recommend to future studies that are planning to use a similar simulation approach that they should work with a larger number of real sampling locations, covering a defined environmental gradient, right from the start to avoid a usage of “fake” integers and to potentially reach a higher predictive power for more microbial taxa in the model. We established our precise applied HMSC modelling framework through multiple test runs, in which we checked model performance indicated by the coefficient of discrimination Tjur’s R^2^ (^26^), which is defined as the difference between the average model prediction for successes and failures and was summarized as the mean Tjur’s R^2^ across species. A two-fold cross-validation was performed to assess the predictive power of the model. Our final model was a HMSC which included a spatial random effect, temperature as only covariate, and that is based on a Poisson distribution. We used HMSC ^27^ from the “Hmsc” R-package ^28^ to fit a joint species distribution model on the key microbial community member level. Our response matrix were the relative microbial abundances of key microbes (transformed to fake integers). We included a random effect at the level of sampling station using a latent factor approach ^29^. After fitting the model (assuming a linear effect), we examined and ensured the convergence of the Markov chain Monte Carlo simulations and evaluated the model fit (psrf values of the beta parameters were on average 1.002 with a 95 % confidence interval= 0.003). The explanatory power of the model was assessed by computing the R^2^ for each microbial species, and as mean across all microbial species. Furthermore, a two-fold cross-validation was conducted to evaluate the predictive power of the model. Subsequently, the parameter estimates were explored and predictions made, followed by an additional step to make spatial predictions. We then standardized or spatial predictions to 100 %, converting “fake counts” into relative abundances. In the next step, we plotted our predictions across the complete temperature range, covered by the subsetted temperature grid. Based on these plots we determined temperature ranges which may have a too high uncertainty of predictability (esp. those temperature ranges that were far of values with actual in situ measurements of microbial community composition). Following these evaluations, we subsetted our initial input grid to a narrower temperature range, by removing all *Vazella pourtalessii* occurrence points above or below the temperature range 6.2-8.2°C. We then reran the complete HMSC model again including all previous steps ((i) set up a HMSC model with a spatial random effect and temperature as only covariate, based on a Poisson distribution, (ii) fit model, (iii) evaluate convergence, (iv) compute and show model fit, (v) show parameter estimates, (vi) make predictions, (vii) make spatial predictions, and standardize them to 100 %. After these steps, we extracted the ratio of explanatory vs predictive power of all 15 key ASVs (**Supplementary Table 3**). We then extracted the two ASVs with the lowest positive values (ratio of 1.2 and 1.9) and plotted the relationship with temperature of those ASVs (temperature gradient plot **Figure 2A**). In the next step we generated individual ASV distribution maps of those two ASVs in space. Furthermore, we constructed a (sparcc-) co-occurrence network based on the total microbial community in *Vazella pourtalesii*. Within Gephi (version 0.9) the Furchterman Reingold algorithm was used (**Supplementary Figure 3A**), from there the graph object was exported to Cytoscape (version 3.10.0) and there projected into a circular layout. For the visualization only those ASVs were visualized that had a connection to at least one key ASV. “Highly interconnected “ and “interconnected” ASVs were defined based on the degree of connectivity inside the network (see explanation above). ASV taxonomic identities were traced back manually. A potential correlation between network connectedness (number of edges) and model fit (ratio of explanatory power vs predictive power) was evaluated through Pearson correlation for all 15 key ASVs.

### Trait predictions of the microbial abundance status in deep-sea sponges in space

In order to perform trait predictions of the microbial abundance status in deep-sea sponges in space, we focused on the Flemish Cap area. For this approach we integrated published data of geographic sponge species distribution ^30^, and knowledge of the microbial abundance status of the different sponge species (^11,31^ for ten sponge species (five HMA species and five LMA species; **Supplementary Table I**). Here, we considered the microbial abundance status (HMA vs LMA) as a binary trait (0 absence in the grid cell, 1 presence in the grid cell), and predicted both, HMA and LMA distributions, on a continuous spatial scale with the help of two separate RF models. BNAM environmental data layers of the present timeframe, as well as slope and depth, that were snapped to the raster extend used in Murillo et al., 2020, were used as environmental data layers. The RF models were run using a similar approach as described above, in brief: (i) fitting of models with the *randomForest*-package in R ^19^ with default parameters and 500 trees, (ii) derivation of model accuracy metrics and threshold probabilities using a 5-fold spatial block cross-validation, where for each cross-validation run the threshold-indepented area under the receiver operating characteristic curve (AUC) was calculated, and the mean standard deviation were derived, (iii) extraction of the MSS, with the help of the *optimal*.*thresholds*-function in the “PresenceAbsence”-R package ^20^, and deriving sensitivity, specificity, and the true skill statistic (TSS; ^21^) from the confusion matrix.

We then correlated our spatial predictions of sponge microbial abundance status occurrence with predictions of overall ecosystem function, i.e. here nutrient cycling and habitat provision (derived from ^30^). For this, the original rasters (created by ^30^) were resampled to the 0.088 cell size using Bilinear interpolation as the resampling technique in ArcGIS Pro (*doi:10*.*17632/fx3vd2tgcf*.*1 (“Data6”)*). Afterwards, a correlation assessment between predictions of sponge microbial abundance occurrence and predictions of overall ecosystem function was done in form of Spearman correlations.

Overall biomasses of each sponge microbial abundance status were calculated, summing up the biomass values of all sponge species (published in ^30^) for each of the two categories. We then computed a biomass network, integrating our newly generated information on HMA and LMA biomasses with biomass measurements of other sessile filter feeding invertebrates, which occur in high abundances at the Flemish Cap. This biomass network represents 116 different species in total, which are combined into functional (passive and active filter feeders) and taxonomic (8 phyla) groups (*doi:10*.*17632/fx3vd2tgcf*.*1 (“Data7”)*). The data in our present paper is a subset of all 285 organisms covered in Murillo et al., 2020, based on only those organisms which fulfilled the criteria: (i) “motility” (= sessile or sessile-burrow), “degree of contagion” (= high abundance or patchy), and “feeding mode” (= active or passive filter feeding). Biomass networks (indicating correlations between biomass values of individual size groups per taxon) were calculated in R and visualised in Cytoscape (version 3.10.0).

## References main manuscript text

1. Roberts, E. et al. Water masses constrain the distribution of deep-sea sponges in the North Atlantic Ocean and Nordic Seas. Mar. Ecol. Prog. Ser.; 2021, 659, 75–96.

2. Beazley, L. et al. Climate change winner in the deep sea? Predicting the impacts of climate change on the distribution of the glass sponge Vazella pourtalesii. Mar. Ecol. Prog. Ser.; 2021, 657, 1–23.

3. Taranto, G. H. et al. Spatial distributions, environmental drivers and co-existence patterns of key cold-water corals in the deep sea of the Azores (NE Atlantic). Deep. Res. Part I Oceanogr. Res. Pap.; 2023, 197.

4. Gasbarro, R., Sowers, D., Margolin, A. & Cordes, E. E. Distribution and predicted climatic refugia for a reef-building cold-water coral on the southeast US margin. Glob. Chang. Biol.; 2022, 28, 7108–7125.

5. Auscavitch, S. R. et al. Distribution of deep-water scleractinian and stylasterid corals across abiotic environmental gradients on three seamounts in the Anegada Passage. PeerJ; 2020, 8, 1–25.

6. Dorey, N., Gjelsvik, Ø., Kutti, T. & Büscher, J. V. Broad thermal tolerance in the cold-water coral Lophelia pertusa from Arctic and boreal reefs. Front. Physiol.; 2020, 10, 1–12.

7. Wang, S., Murillo, F. J. & Kenchington, E. Climate-change refugia for the bubblegum coral Paragorgia arborea in the Northwest Atlantic. Front. Mar. Sci.; 2022, 9, 1–23.

8. Davies, A. J., Wisshak, M., Orr, J. C. & Murray Roberts, J. Predicting suitable habitat for the cold-water coral Lophelia pertusa (Scleractinia). Deep. Res. Part I Oceanogr. Res. Pap.; 2008, 55, 1048–1062.

9. Morato, T. et al. Climate-induced changes in the suitable habitat of cold-water corals and commercially important deep-sea fishes in the North Atlantic. Glob. Chang. Biol.; 2020, 26, 2181–2202.

10. Busch, K. et al. Biodiversity, environmental drivers, and sustainability of the global deep-sea sponge microbiome. Nat. Commun.; 2022, 13.

## References Methods

1. R Development Core Team. R: A language and environment for statistical computing. https://doi.org/http://www.r-project.org. 2008.

2. ESRI. ArcGIS Desktop: Release 10. 2011.

3. van Rossum, G. Python tutorial, Technical Report CS-R9526. Centrum voor Wiskunde en Informatica (CWI). 1995.

4. Beazley, L. et al. Climate change winner in the deep sea? Predicting the impacts of climate change on the distribution of the glass sponge Vazella pourtalesii. Mar. Ecol. Prog. Ser. 2021; 657, 1–23.

5. GEBCO Bathymetric Compilation Group. The GEBCO_2019 Grid - a continuous terrain model of the global oceans and land. 2019.

6. Wang, Z., Lu, Y., Greenan, B., Brickman, D. & Detracey, B. BNAM : An eddy-resolving North Atlantic Ocean model to support ocean monitoring. Can. Tech. Rep. Hydrogr. Ocean Sci. 2018; 327, vii + 18p.

7. Carton, J. A. & Giese, B. S. A reanalysis of ocean climate using Simple Ocean Data Assimilation (SODA). Mon. Weather Rev. 2008; 136, 2999–3017.

8. IPCC. Climate change 2013: the physical science basis. Contribution of Working Group I to the Fifth Assessment Report of the Intergovernmental Panel on Climate Change. Cambridge University Press. 2013.

9. Brickman, D., Wang, Z. & Detracey, B. High resolution future climate ocean model simulations for the Northwest Atlantic Shelf region. Canadian Technicial Report of Hydrography and Ocean Sciences. 2016; 315.

10. SODA. https://coastwatch.pfeg.noaa.gov/erddap/griddap/hawaii_d90f_20ee_c4cb_LonPM180.html. 2023.

11. Busch, K. et al. Biodiversity, environmental drivers, and sustainability of the global deep-sea sponge microbiome. Nat. Commun. 2022; 13.

12. Kellogg, C. A. & Pratte, Z. A. Unexpected diversity of Endozoicomonas in deep-sea corals. Mar. Ecol. Prog. Ser. 2021; 673, 1–15.

13. Morato, T. et al. Climate-induced changes in the suitable habitat of cold-water corals and commercially important deep-sea fishes in the North Atlantic. Glob. Chang. Biol. 2020; 26, 2181–2202.

14. NOAA. NOAA Deep-sea Coral Data Portal. https://deepseacoraldata.noaa.gov. 2023.

15. Murillo, F. J., Serrano, A., Kenchington, E. & Mora, J. Epibenthic assemblages of the Tail of the Grand Bank and Flemish Cap (northwest Atlantic) in relation to environmental parameters and trawling intensity. Deep. Res. Part I Oceanogr. Res. Pap. 2016; 109, 99–122.

16. Murillo, F. J. et al. Sponge assemblages and predicted archetypes in the eastern Canadian Arctic. Mar. Ecol. Prog. Ser. 2018; 597, 115–135.

17. Steffen, K. et al. Oceanographic setting influences the prokaryotic community and metabolome in deep-sea sponges. Sci. Rep. 2022; 12, 1–16.

18. iNaturalist. iNaturalist Database. 2023.

19. Liaw, A. & Wiener, M. Classification and regression by randomForest. R News. 2002; 2, 18–22.

20. Freeman, E. A. & Moisen, G. PresenceAbsence: An R package for PresenceAbsence analysis. J. Stat. Softw. 2008; 23, 1–31.

21. Allouche, O., Tsoar, A. & Kadmon, R. Assessing the accuracy of species distribution models: Prevalence, kappa and the true skill statistic (TSS). J. Appl. Ecol. 2006; 43, 1223–1232.

22. Lopes, P. F. M., Verba, J. T., Begossi, A. & Pennino, M. G. Predicting species distribution from fishers’ local ecological knowledge: A new alternative for data-poor management. Can. J. Fish. Aquat. Sci. 2019; 76, 1423–1431.

23. Bolyen, E. et al. QIIME 2: Reproducible, interactive, scalable and extensible microbiome data science using QIIME 2. Nat. Biotechnology. 2019; 37:852–857.

24. Callahan, B. J. et al. DADA2: High-resolution sample inference from Illumina amplicon data. Nat. Methods. 2016; 13, 581–583.

25. Quast, C. et al. The SILVA ribosomal RNA gene database project: Improved data processing and web-based tools. Nucleic Acids Res. 2013; 41, 590–596.

26. Tjur, T. Coefficients of determination in logistic regression models - A new proposal: The coefficient of discrimination. Am. Stat. 2009; 63, 366–372.

27. Ovaskainen, O. et al. How to make more out of community data? A conceptual framework and its implementation as models and software. Ecol. Lett. 2017; 20, 561–576.

28. Tikhonov, G., Opedal, Ø.H., Abrego, N., Lehikoinen, A., de Jonge, M.M.J., Oksanen, J., Ovaskainen, O. Joint species distribution modelling with the r-package Hmsc. Methods Ecol. Evol. 2019; 11, 442–447.

29. Ovaskainen, O., Abrego, N., Halme, P. & Dunson, D. Using latent variable models to identify large networks of species-to-species associations at different spatial scales. Methods Ecol. Evol. 2016; 7, 549–555.

30. Murillo, F. J., Weigel, B., Bouchard Marmen, M. & Kenchington, E. Marine epibenthic functional diversity on Flemish Cap (north-west Atlantic)—Identifying trait responses to the environment and mapping ecosystem functions. Divers. Distrib. 2020; 26, 460–478.

31. Maldonado, M., Bayer, K. & López-Acosta, M. Nitrogen and phosphorus cycling through marine sponges: Physiology, cytology, genomics, and ecological implications. Frontiers in Invertebrate Physiology: A Collection of Reviews. Apple Academic Press. 2024.

