## Supplementary Material for "Past, present, and future spatial distributions of deep-sea coral and sponge microbiomes revealed by predictive models"

to:

contains:

- Supplementary Figures 1-8 & I-II
- Supplementary Tables 1-3 & I-III
- Supplementary Texts 1-3

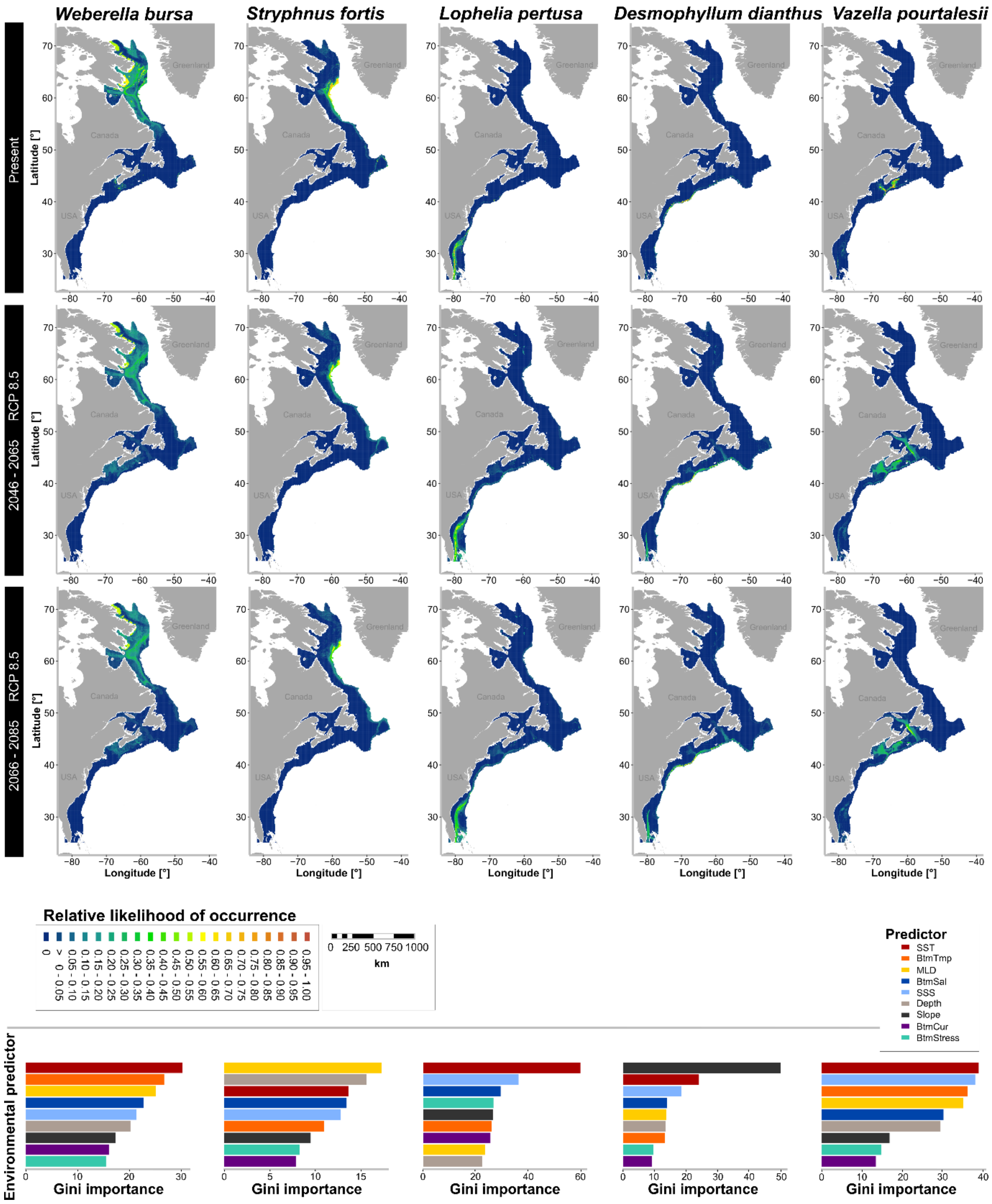


**Supplementary Figure 1** Relative likelihood of occurrence of *Weberella bursa*, *Stryphnus fortis*, *Lophelia pertusa*, *Desmophyllum dianthus*, and *Vazella pourtalesii* from Random Forest predictions under present day and future (RCP8.5 2046-2065 and 2066-2085) climatic conditions. Gini importances (bar charts) indicate ranking of environmental predictor importance in Random Forest models for each animal species.

**Supplementary Table 1** Accuracy measures for Random Forest models trained and tested on the *Weberella bursa, Stryphnus fortis, Lophelia pertusa, Desmophyllum dianthus*, and *Vazella pourtalesii* presence/pseudo-absence data of our study area. Cross-validation was done via 5-fold spatial blocking with random assignment of blocks into folds. Sensitivity, specificity, and the true skill statistic (TSS) were generated from a confusion matrix of tabulated outcomes that was thresholded using the maximum of sensitivity + specificity (MSS) identified each model. AUC: area under the receiver operating characteristic curve.

| **Animal species** | **Mean AUC + - SD** | **Sensitivity + - SD** | **Specificity + - SD** | **TSS** | **MSS threshold** |
|---|---|---|---|---|---|
| *Weberella bursa* | 0.83 +- 0.01 | 0.63 +- 0.05 | 0.87 +- 0 | 0.49 | 0.03 |
| *Stryphnus fortis* | 0.96 +- 0.01 | 0.93 +- 0.04 | 0.90 +- 0 | 0.83 | 0.02 |
| *Lophelia pertusa* | 0.95 +- 0.01 | 0.94 +- 0.02 | 0.89 +- 0 | 0.83 | 0.02 |
| *Desmophyllum dianthus* | 0.94 +- 0.00 | 0.92 +- 0.03 | 0.86 +- 0 | 0.78 | 0.01 |
| *Vazella pourtalesii* | 0.96 +- 0.01 | 0.95 +- 0.02 | 0.85 +- 0 | 0.80 | 0.02 |

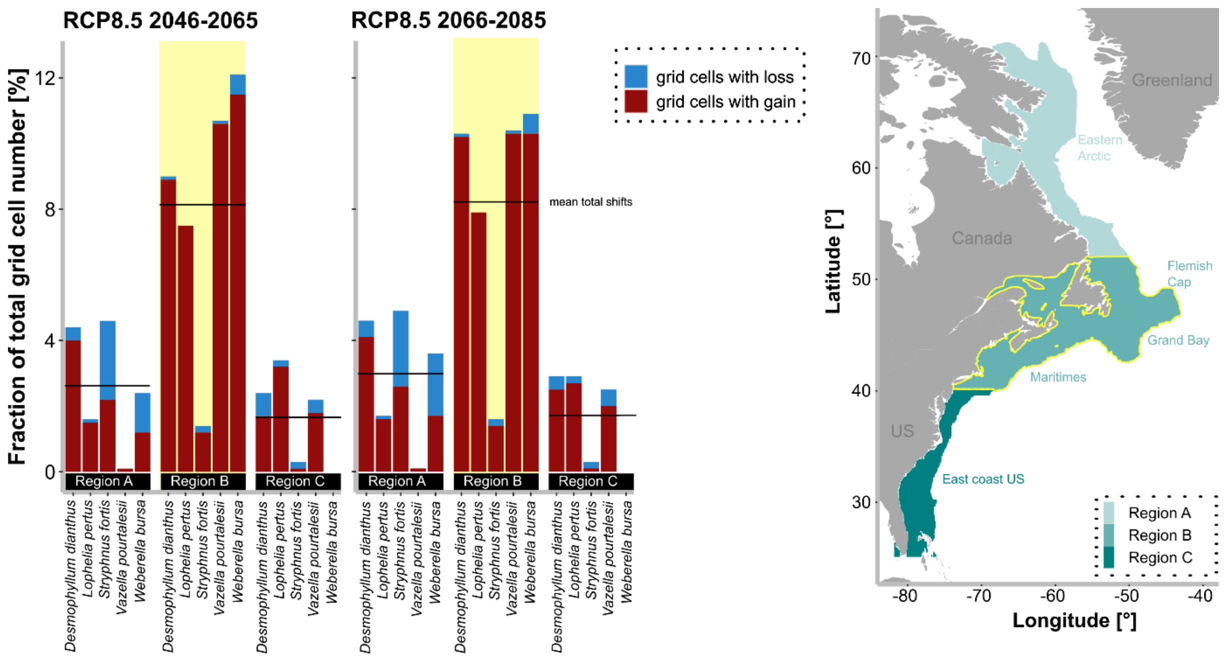


**Supplementary Figure 2** Predicted future shifts in cumulative microbial richness in deep-sea corals and sponges across the three sub-regions (A, B, and C) indicated for the five coral and sponge species, given as fraction of total grid cell number. Mean total shifts are indicated by lines. As sub-region B depicts the largest shifts, it is highlighted in yellow within the barplot, but also on the map next to it.

**Supplementary Text 1:** *Some additional information on the identified key ASVs*

We identified 15 key microbial taxa (see **Methods** for criteria on how they were identified) and we consider them to be representative of the overall *Vazella pourtalesii* microbial community as (i) they all together cover the majority (i.e. 67 %) of the total *Vazella pourtalesii* microbial community in terms of relative abundances, and (ii) their taxonomic composition, also strongly resembles the community composition described in [1]. Out of these 15 key microbial taxa we identified two key microbial taxa which had a modelable relationship with temperature when using our model. Taxonomically, one of the two microbial taxa belonged to the phylum Patescibacteria (class Parcubacteria, order Candidatus Kaiserbacteria), while the other microbial taxon remained unclassified. Our previous work ([1]) aimed to identify functional strategies and metabolic networks of the main *Vazella pourtalesii* symbionts. Patescibacteria were identified as anaerobes with reduced genomes that tap into the microbial community for resources ([1]).

**Supplementary Text 2:** *Some additional information on the generated microbial co-occurrence networks*

In order to evaluate how shifts in relative abundances of the key temperature-driven ASVs translate into shifts in overall microbial community composition, we constructed co-occurrence networks (**Supplementary Figure 3**). Those co-occurrence networks overall resembled the strong intertwining of *Vazella pourtalesiis* key microbes: Seven of the 15 key ASVs showed network connections to a large number of additional other ASVs (and were therefore called "highly interconnected ASVs"), while eight ASVs had network connections to solely other key ASVs (and were therefore called "interconnected ASVs") (**Supplementary Figure 3A**). The number of positive and negative correlations was more or less balanced for the complete network, and the two key temperature-driven ASVs split across both groups (highly interconnected ASVs and interconnected ASVs) (**Supplementary Figure 3B**). This suggests that changes in relative abundances of ASVs in relation to changes in temperature may translate into cascading changes in overall microbial community composition. We initially hypothesized that the HMSC-model fit of temperature-dependence would decrease for ASVs with a large number of network connections. We came to this hypothesis because we would expect a "biotic buffering effect" against changing abiotic conditions upon a stronger connectedness between microbial taxa. The number of network connections and the model ratio (explanatory power vs predictive power) were however not significantly correlated (Pearson correlation -0.007, p-value=0.980), which suggests that changes in abiotic conditions may not only impact single microbial taxa, but indeed a larger network of taxa. Furthermore, the taxonomic composition of both, highly interconnected and interconnected ASVs were composed of similar taxonomic groups (Proteobacteria, Patescibacteria, and unclassified).

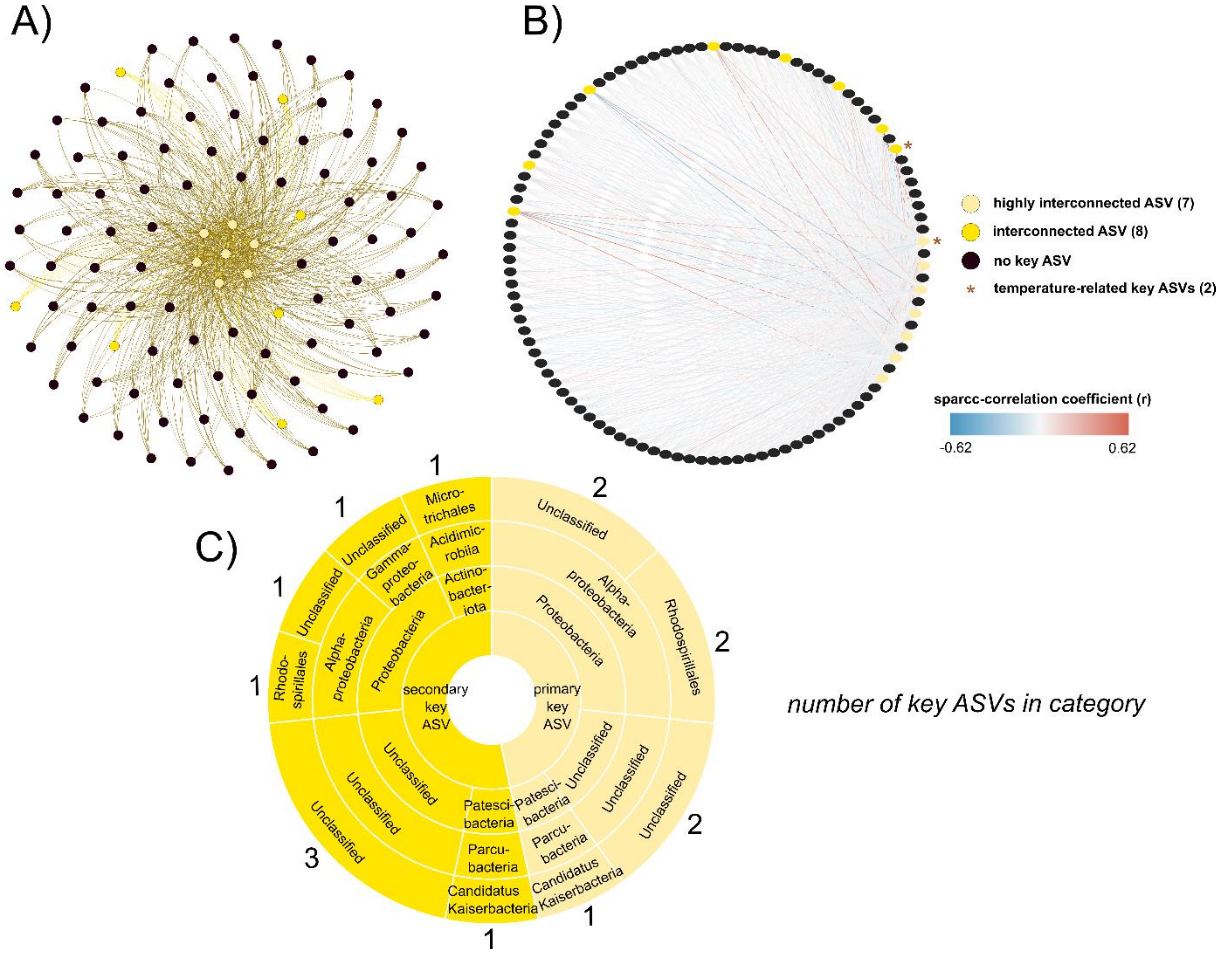


**Supplementary Figure 3 A)** Furchterman Reingold network presentation, highlighting seven key ASVs with particularly high network connections in the central part of the network. Those seven key ASVs are called “highly interconnected ASVs” in the following. Other key ASVs which have less network connections (and only to other key ASVs) are also indicated by yellow color, and called “interconnected ASVs” in the following. ASVs which have a connection with a key ASV, but are not key ASVs themselves, are marked by black color. **B)** Circular network presentation of the same network as in A). Directions of correlations between individual ASVs are marked by colors: blue=negative correlation, red=positive correlation. The same yellow and black color code is applied as in A). In addition, temperature-related key ASVs are marked by an asterisk. **C)** Sunburst diagram, showing taxonomic composition of highly interconnected and interconnected ASVs (phylum, class, and order-level). The number of key ASVs in each category is written at the outer part of the plot.

**Supplementary Text 3** *Trait-based approaches*

One particularly interesting trait of sponges is their separation into HMA (high microbial abundance) sponges and LMA (low microbial abundance) sponges. As the name “HMA-LMA dichotomy” suggests, some sponges contain dense microbial consortia in their tissues, while other species lack such dense communities. This status (HMA or LMA) correlates not only with microbial abundance and richness, but also with differences in ecosystem function (e.g. in terms of their nutrient cycling capacity) of both metaorganism types. Following predictions of cumulative microbial richness in corals and sponges, as well as predictions of individual microbial taxa and overall microbial community composition in relation to environmental shifts and gradients, we sought to explore the usability of trait-based modelling approaches. Focusing on the Flemish Cap area, we assigned the HMA-LMA status to ten key sponge species, and used occurrence data of those species to predict the spatial distribution of the HMA-LMA-status at the Flemish Cap (**Supplementary Figure 4 A+B**; **Supplementary Table I**, **Supplementary Table II**). The HMA-status was predicted to occur exclusively at the rim of the Flemish Cap and the main correlating physical parameter with this status is temperature (**Supplementary Figure I**). In contrast, occurrence of the LMA status sponges was predicted to be much more widespread through the Flemish Cap. The main correlating physical parameter of this status is salinity. However, it also has to be noted that the predictive capacity of the model was worse for LMA sponges than for HMA sponges, which may be related to a larger intra-group variability of LMA sponges. Interestingly we found a slight correlation between occurrence of the HMA status and overall predicted nutrient cycling capacity, and a stronger one between occurrence of the LMA status and overall predicted nutrient cycling capacity (both significant with p-value<0.001). A similar trend was observed for a correlation between each sponge microbial abundance status and overall predicted habitat provision (**Supplementary Table III**). Interestingly, the clear spatial distribution pattern of HMA-occurrence seems to be driven strongly by presence of geodids, and comparisons with historic data (dating back to >16.7 ka BP) suggest that this pattern has most likely existed at the Flemish Cap for a long time (**Supplementary Figure II**). Interestingly, although the HMA status is predicted to occur at a much narrower range in the slope waters at the Flemish Cap, biomass per location is much higher for HMA sponges than for LMA ones (**Supplementary Figure 4C**). A biomass network with other filter-feeding invertebrates at the Flemish Cap suggests that HMA sponge biomass is mainly correlated with cnidarians (corals) and other sponges, while biomass of LMA sponges is correlated with a much larger number of taxa (**Supplementary Figure 4D**). We believe that evaluating microbial-macrobial relationships through such a trait-based approach represents a promising tool to further incorporate microbial data into a larger ecosystem context and into larger foodweb analyses. It may be particularly interesting to evaluate the role of the microbiome in facilitating the host to function as a capable space competitor (see also literature on space competition between sponges and corals, e.g. [2]).

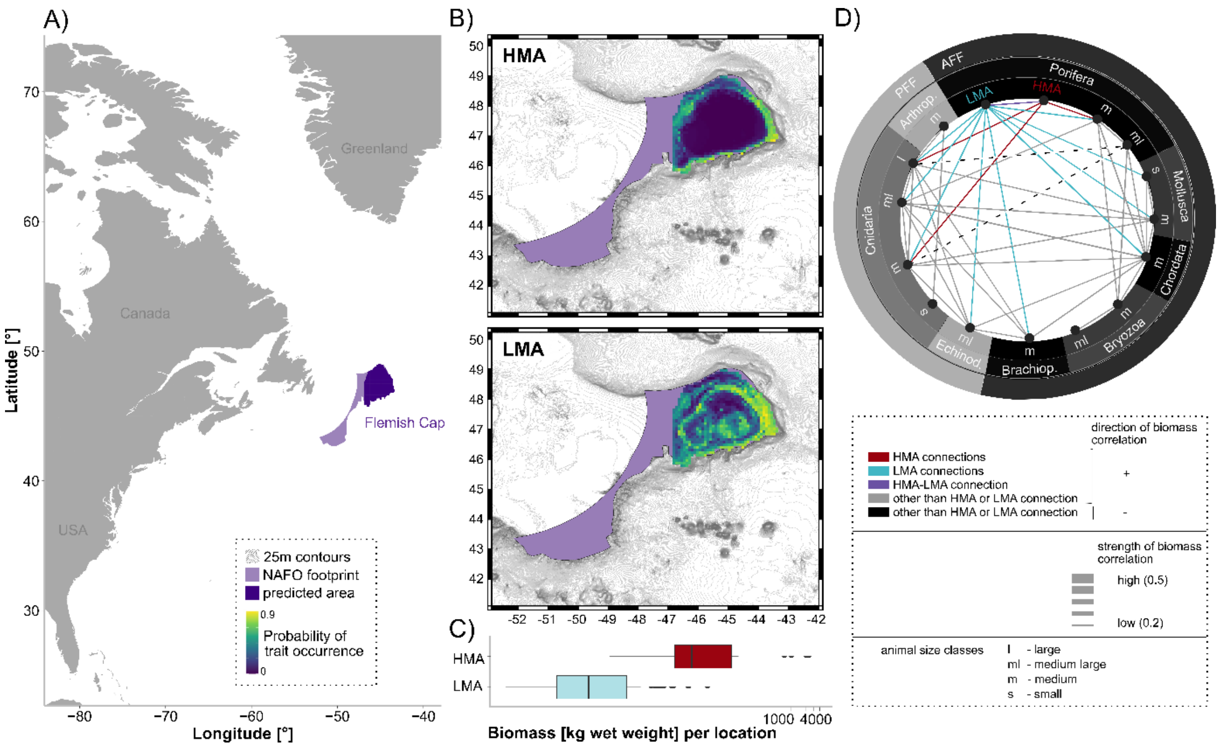


**Supplementary Figure 4 A)** Overview of study area location. The Flemish Cap is indicated by dark purple colour. **B)** RF-probability maps of HMA- and LMA-occurrence. **C)** Comparison of biomass per location for HMA against LMA sponge species. **D)** Biomass-correlation-network between HMA- and/or LMA-sponges with other filter feeding invertebrates, which occur in high abundances at the Flemish Cap. The biomass network represents 116 different species in total, which are combined into two functional groups (PFF:passive and AFF:active filter feeders), eight phyla (Porifera, Mollusca, Chordata, Bryozoa, Brachiopoda, Echinodermata, Cnidaria, Arthropoda), and four size classes (l=large, ml=medium large, m=medium, s=small). Connections of taxa with HMA sponges are colored in red, connections with LMA sponges in blue, and connections between HMA and LMA sponges in purple. Other positive correlations are colored in grey, while the only two negative connections (between medium large Porifera and medium Cnidaria, as well as large Cnidaria, respectively) are indicated by dashed lines and a black colour.

**Supplementary Table I** Overview of microbial abundance status of key sponge species at the Flemish Cap. A-B= asexual-budding, BS=sexual broadcast spawner, BID=sexual brooder-indirect development pelagic, BDD=sexual brooder-direct development demersal or viviparous, PP=pelagic planktotrophic, PL= pelagic lecithotrophic, B= benthic.

| **Taxa** | **Status** | **Longevity** | **Reproductive method** | **Propagule dispersal** |
|---|---|---|---|---|
| *Asconema foliatum* | LMA | > 50 years | BID | PL |
| *Geodia barretti* | HMA | > 50 years | A-B, BS | PL |
| *Geodia macandrewii* | HMA | > 50 years | A-B, BS | PL |
| *Geodia parva-phlegraei* | HMA | > 50 years | A-B, BS | PL |
| *Mycale (Mycale) lingua* | LMA | > 50 years | A-B, BID | PL |
| *Stelletta normani* | HMA | > 50 years | A-B, BS | PL |
| *Stryphnus fortis* | HMA | > 50 years | A-B, BS | PL |
| *Stylocordyla borealis* | LMA | > 50 years | BDD | B |
| *Tentorium semisuberites* | LMA | > 50 years | A-B, BS | PL |
| *Weberella bursa* | LMA | > 50 years | A-B, BS | PL |

**Supplementary Table II** Accuracy measures for Random Forest models trained and tested on the HMA and LMA data of the Flemish Cap. Cross-validation was done via 5-fold spatial blocking with random assignment of blocks into folds. Sensitivity, specificity, and the true skill statistic (TSS) were generated from a confusion matrix of tabulated outcomes that was thresholded using the maximum of sensitivity + specificity (MSS) identified each model. AUC: area under the receiver operating characteristic curve.

| **Host microbial abundance status** | **Mean AUC + - SD** | **Sensitivity + - SD** | **Specificity + - SD** | **TSS** | **MSS threshold** |
|---|---|---|---|---|---|
| HMA | 0.76 +- 0.09 | 1.00 +- 0.00 | 0.60 +- 0.04 | 0.60 | 0.05 |
| LMA | 0.58 +- 0.03 | 0.64 +- 0.06 | 0.53 +- 0.05 | 0.17 | 0.44 |

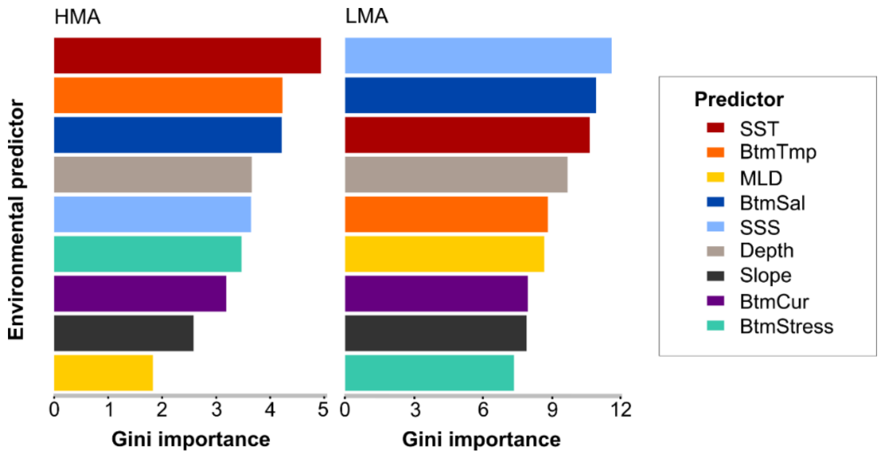


**Supplementary Figure I** Gini importances indicate ranking of environmental predictor importance in Random Forest models for each microbial abundance status.

**Supplementary Table III** Results of Spearman correlations between predictions of two overall ecosystem functions at the Flemish Cap (taken from [3]) and spatial predictions of sponge microbial abundance status occurrence.

| | Nutrient cycling | | Habitat provision | |
|---|---|---|---|---|
| **Group** | **Correlation coefficient** | **p-value** | **Correlation coefficient** | **p-value** |
| HMA | 0.217 | <0.001 | 0.463 | <0.001 |
| LMA | 0.665 | <0.001 | 0.589 | <0.001 |

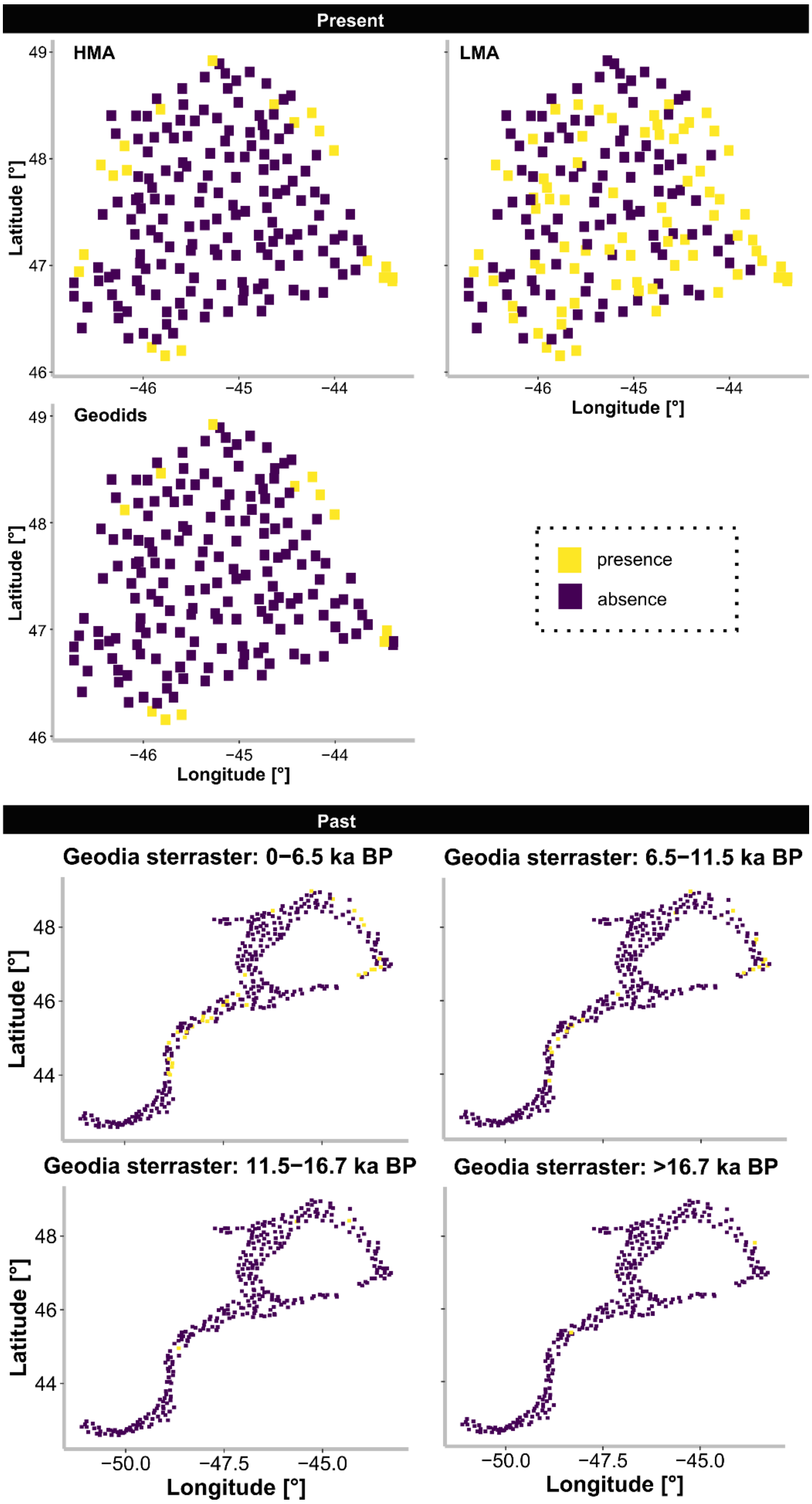


**Supplementary Figure II** Trait over time. **Upper panel:** Present raw occurrence data of the HMA-status, the LMA-status, and geodiids at the Flemish Cap. Data was taken from [3]. **Lower panel:** Records of *Geodia* sterrasters found in different age layers in cores taken at the Flemish Cap. Data was taken from[4]. Note that presence of sterrasters occurs at similar places from the past to the present.

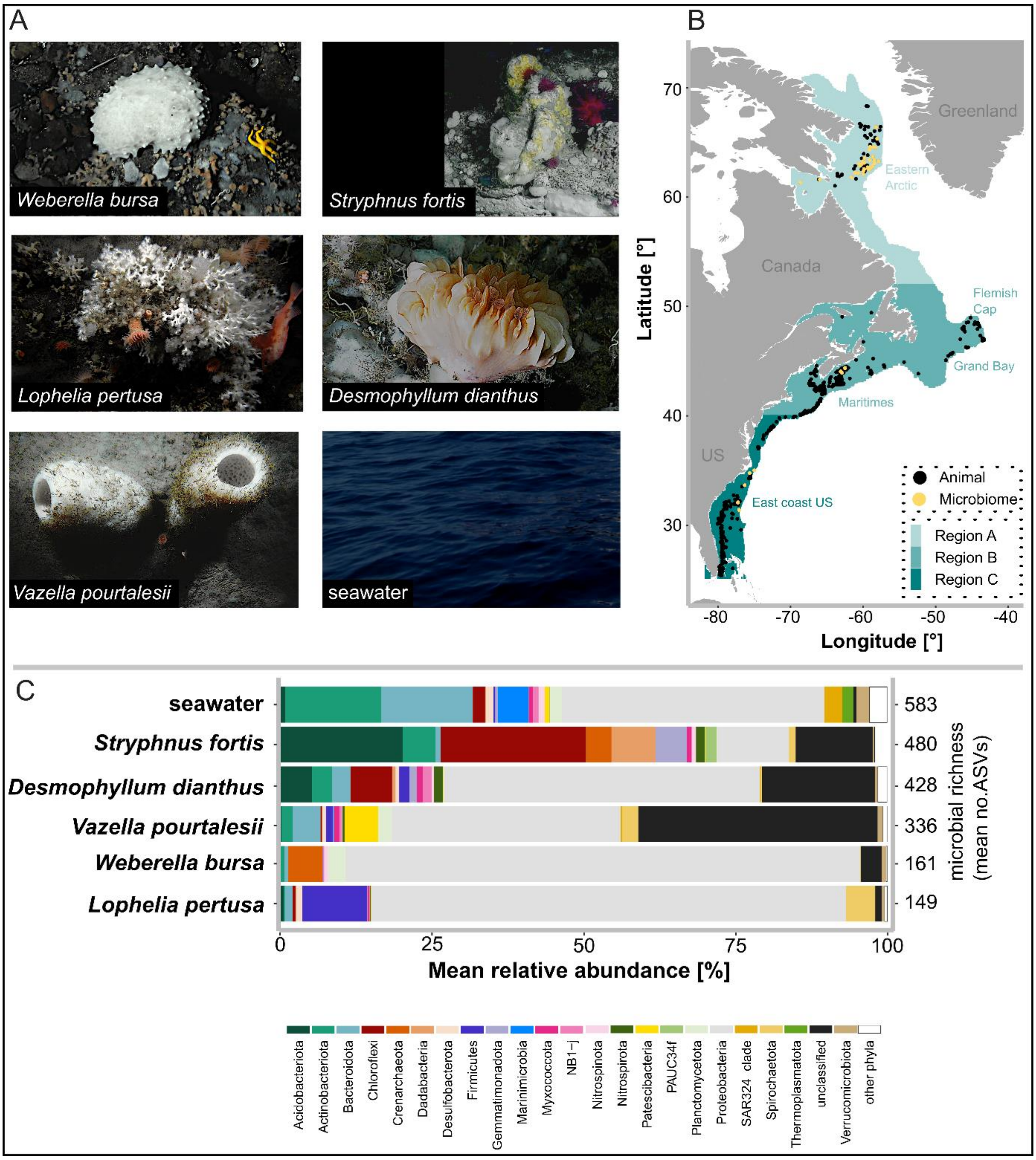


**Supplementary Figure 5 A)** Underwater photographs of study animal species *Weberella bursa*, *Stryphnus fortis*, *Lophelia pertusa*, *Desmophyllum dianthus*, and *Vazella pourtalesii*, as well as seawater reference. **B)** Overview of study area, divided into three regions: Region A, Region B, and Region C. Animal and microbe occurrences used for analyses of this study are indicated by black and yellow dots, respectively. **C)** Mean relative abundances of microbial phyla per sample type. Sample types are sorted after mean microbial richness, in descending order from top to bottom. Unclassified microbial taxa are marked by black bars.

| | Time frame | Environmental dataset |
|---|---|---|
| Past | 1871-1900 | SODA |
| | 1901-1930 | SODA |
| | 1931-1960 | SODA |
| | 1961-1989 | SODA |
| Present | 1990-2015 | BNAM |
| Future | 2046-2065 | BNAM (RCP8.5) |
| | 2066-2085 | BNAM (RCP8.5) |

**Supplementary Figure 6** Overview of analysed time frames and environmental datasets used in this study.

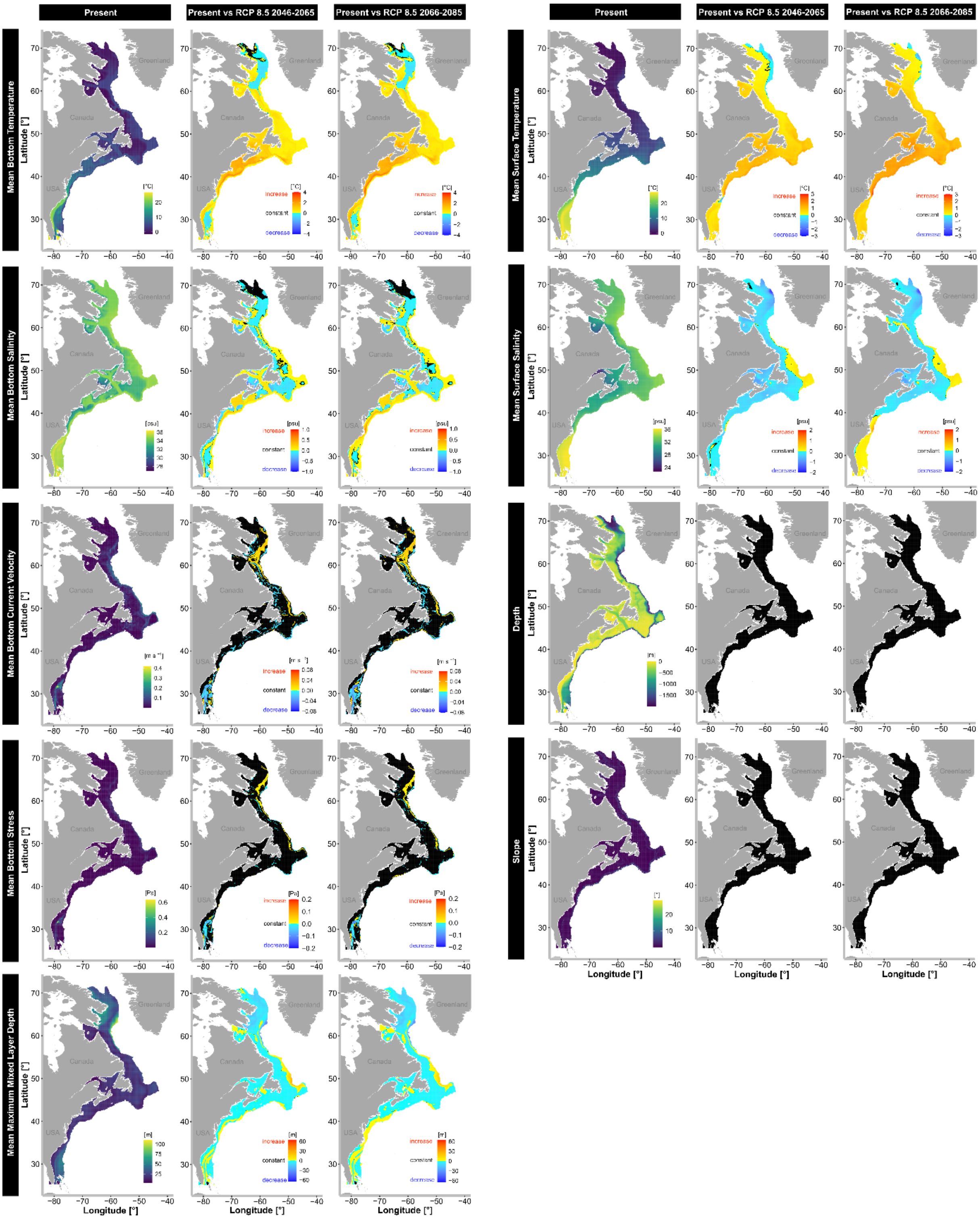


**Supplementary Figure 7** Nine environmental parameters used for species distribution modelling for the present and two future timeframes. Oceanographic parameters were derived from BNAM. Maps for the present are given as absolute values, while the two future timeframes are composed delta-vales, showing differences in comparison to the present.

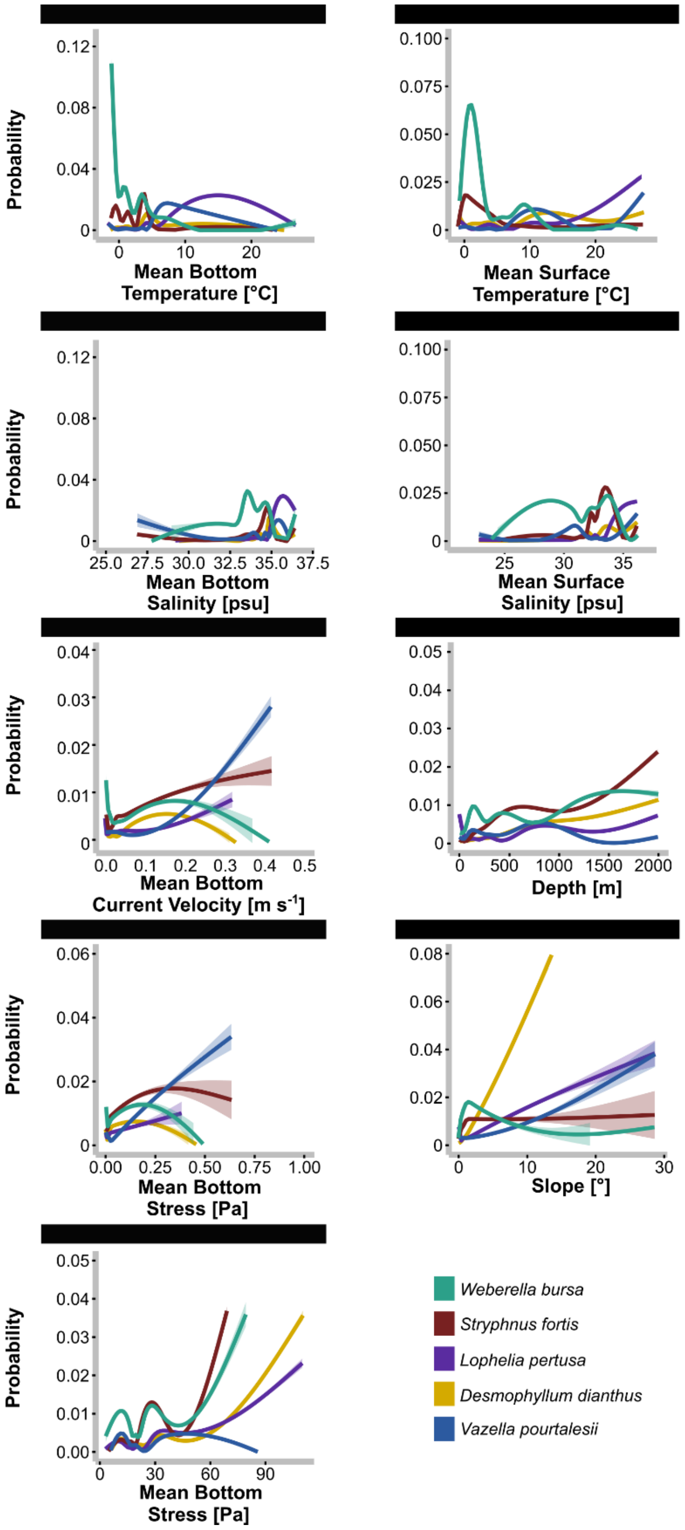


**Supplementary Figure 8** Functional response curves, showing probabilities of animal occurrence for the five different species *Weberella bursa* (green), *Stryphnus fortis* (red), *Lophelia pertusa* (purple*),* *Desmophyllum dianthus* (yellow), and *Vazella pourtalesii* (blue) at different magnitudes of the nine evaluated environmental parameters. Approximate 95% confidence intervals are indicated by ribbons.

**Supplementary Table 2** Summed predicted occupied area [km²] per host species across the seven different time frames.

| **Year** | ***Desmophyllum dianthus*** | ***Lophelia pertusa*** | ***Stryphnus fortis*** | ***Vazella pourtalesii*** | ***Weberella bursa*** |
|---|---|---|---|---|---|
| 1871 - 1900 | 1,926,005 | 1,855,677 | 636,157 | 945,122 | 1,834,066 |
| 1901 - 1930 | 1,912,042 | 1,827,308 | 632,242 | 897,097 | 1,837,189 |
| 1931 - 1960 | 1,872,229 | 1,797,215 | 655,113 | 901,840 | 1,799,432 |
| 1961 - 1989 | 1,876,922 | 1,791,341 | 655,889 | 891,873 | 1,810,119 |
| 1990 - 2015 | 252,847 | 217,412 | 312,258 | 175,161 | 562,537 |
| 2046 - 2065 | 550,787 | 518,704 | 327,759 | 474,057 | 823,030 |
| 2066 - 2085 | 615,282 | 512,827 | 335,448 | 469,983 | 778,308 |

**Supplementary Table 3** Overview of HMSC-model performance (explanatory power, predictive power, and ratio of explanatory power vs predictive power), as well as taxonomic names of the 15 key microbial taxa. Sometimes “bacteria” is abbreviated by “bac.”, and “Candidatus” by “C.”

| Name of ASV | Explana-tory | Predic-tive | Ratio | Phylum | Class | Order |
|---|---|---|---|---|---|---|
| de44846a82b698f04dcc9cba350cbb8b | 0.806 | 0.001 | 850.5 | Proteobac. | Alphaproteobac. | |
| X88e0bc305eefd90b66bc9df365cd2f86 | 0.806 | 0.001 | 850.5 | | | |
| X5fd878722e2825a002fb3459998d2f74 | 0.808 | 0.002 | 406.2 | Proteobac. | Gammaproteobac. | |
| f25abba54a1a8042cfee90caae9eaac2 | 0.770 | 0.004 | 192.9 | Proteobac. | Alphaproteobac. | Rhodospirillales |
| X4928b278e61381feca2dbec26ce8aa6c | 0.770 | 0.017 | 45.4 | Proteobac. | Alphaproteobac. | Rhodospirillales |
| a55e6860b2015dc7390a8e336e772f11 | 0.808 | 0.020 | 40.4 | Proteobac. | Alphaproteobac. | |
| c2de3f4111059c39c137fc464ac1b60c | 0.806 | 0.038 | 21.4 | Proteobac. | Alphaproteobac. | Rhodospirillales |
| d62803e4b3ddeb077fafa7235cff329d | 0.727 | 0.185 | 3.9 | Patescibac. | Parcubacteria | C. Kaiserbacteria |
| X0b3da0a4e278462ae0fbc0c57572d068 | 0.771 | 0.218 | 3.5 | | | |
| bcb979ea7d6db9f1e81929d5a490002c | 0.678 | 0.352 | 1.9 | | | |
| X016474f092254f68cbfe8b808fbdebe8 | 0.771 | 0.643 | 1.2 | Patescibac. | Parcubacteria | C. Kaiserbacteria |
| X5087364294032ebd0b1a43a192c67ad9 | 0.572 | -0.205 | -2.8 | | | |
| X4bb9b107c0e510457ce1a865308b76ea | 0.572 | -0.105 | -5.4 | Proteobac. | Alphaproteobac. | |
| a69ce95b29bec5ec5f35e338b1e2d547 | 0.807 | -0.025 | -32.7 | | | |
| X7526df265f1117b77e0ee234926eebe7 | 0.807 | -0.025 | -32.7 | Actinobac. | Acidimicrobiia | Microtrichales |

### References Supplementary Materials


1. Bayer, K. *et al.* Microbial strategies for survival in the glass sponge *Vazella pourtalesii*. *mSystems.* 2020; **5**, 1–20.
2. Aerts, L. A. M. & Van Soest, R. W. M. Quantification of sponge/coral interactions in a physically stressed reef community, NE Colombia. *Mar. Ecol. Prog. Ser.* 1997; **148**, 125–134.
3. Murillo, F. J., Weigel, B., Bouchard Marmen, M. & Kenchington, E. Marine epibenthic functional diversity on Flemish Cap (north-west Atlantic) - Identifying trait responses to the environment and mapping ecosystem functions. *Divers. Distrib.* 2020; **26**, 460–478.
4. Murillo, F. J., Kenchington, E., Lawson, J. M., Li, G. & Piper, D. J. W. Ancient deep-sea sponge grounds on the Flemish Cap and Grand Bank, northwest Atlantic. *Mar. Biol.* 2016; **163**, 1–11.
